## Supplementary Figures for "A tRNA modification in *Mycobacterium tuberculosis* facilitates optimal intracellular growth"

**Affiliations:**

Supplementary Figures

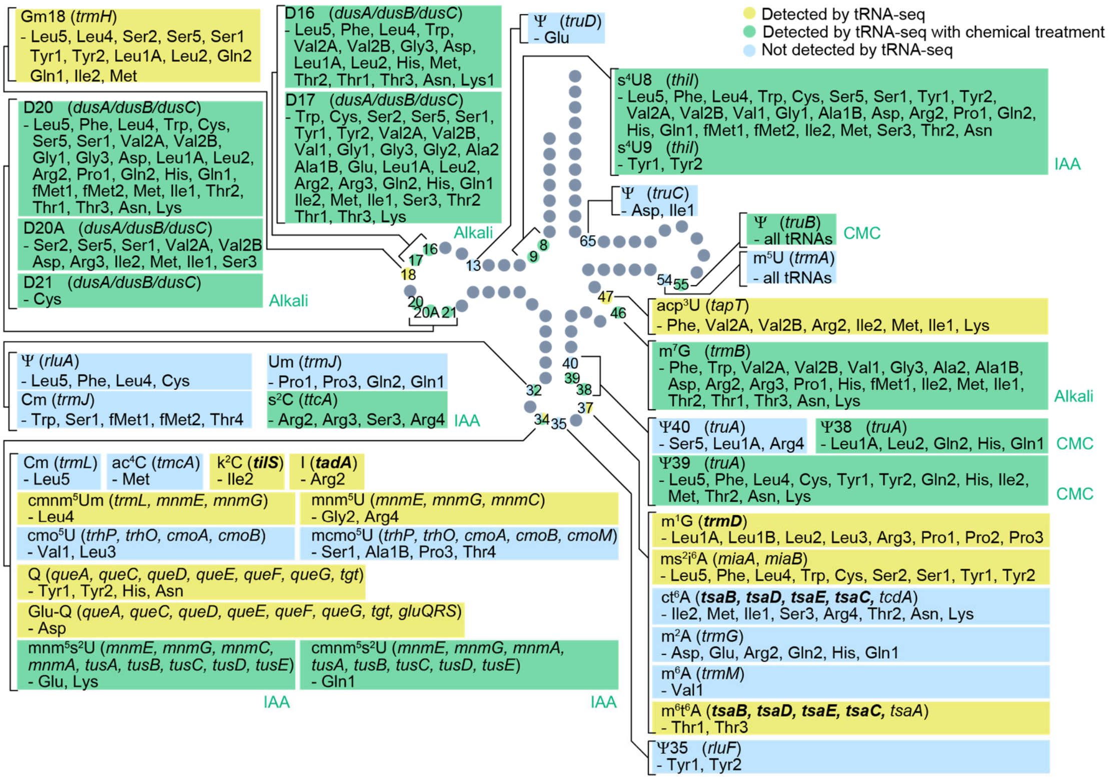


**Supplementary Fig. 1 *E. coli* tRNA modifications detected by tRNA-seq**
Schematic tRNA secondary structure with sites of modifications. Modifications and positions that are detected by RT-derived signature in yellow (without chemical treatment in ref. 14) and green (with chemical treatment in this study), whereas modifications that are not detected by RT-derived signature are shown in light blue. Sites shown in yellow or green were not detected in all the listed tRNA species known to be modified. Genes reported to be essential in *E. coli* are shown in bold.

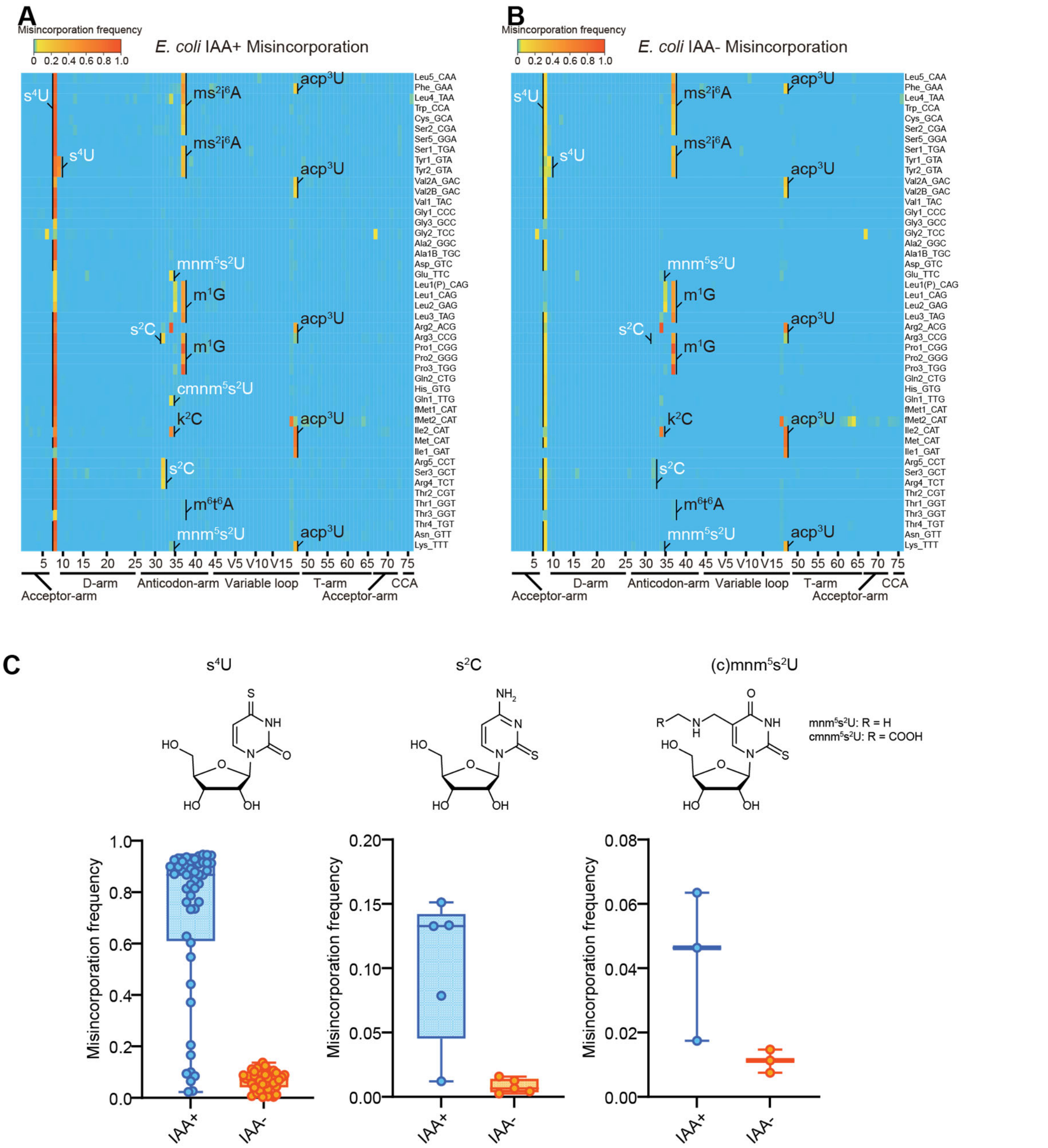


**Supplementary Fig. 2 IAA treatment promotes detecting sulfur modifications by enhancing misincorporation signals.**

(A, B) Heatmaps of the misincorporation signals of *E. coli* tRNAs treated with (A) or without (B) IAA. Known modification sites, including sulfur modifications ($s^4U$, $s^2C$, $s^2U$ in white) are shown.

(C) Misincorporation frequency at $s^4U$, $s^2C$, and $s^2U$ sites of tRNAs treated with or without IAA.

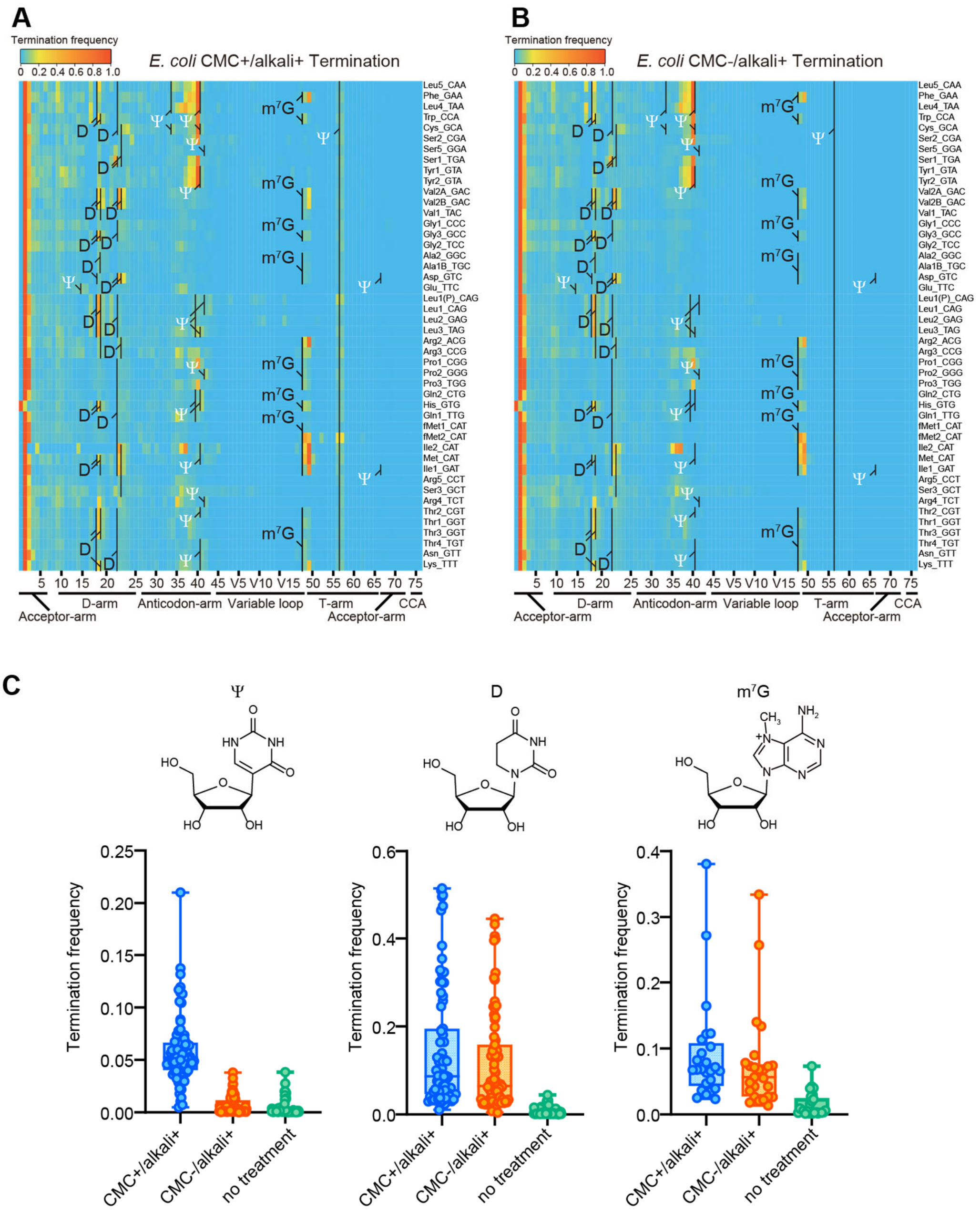


**Supplementary Fig. 3. CMC and the following alkali treatment facilitate detecting additional modifications.**

(A, B) Heatmaps of the termination signals of *E. coli* tRNAs treated with (A) or without (B) CMC. In both conditions, tRNAs are incubated at an alkali condition. Known Ψ (in white), D, $m^7G$ sites are shown.

(C) Termination frequency at known Ψ, D, $m^7G$ sites of tRNAs treated with CMC+/alkali+ or CMC-/alkali+, and tRNAs without treatment.

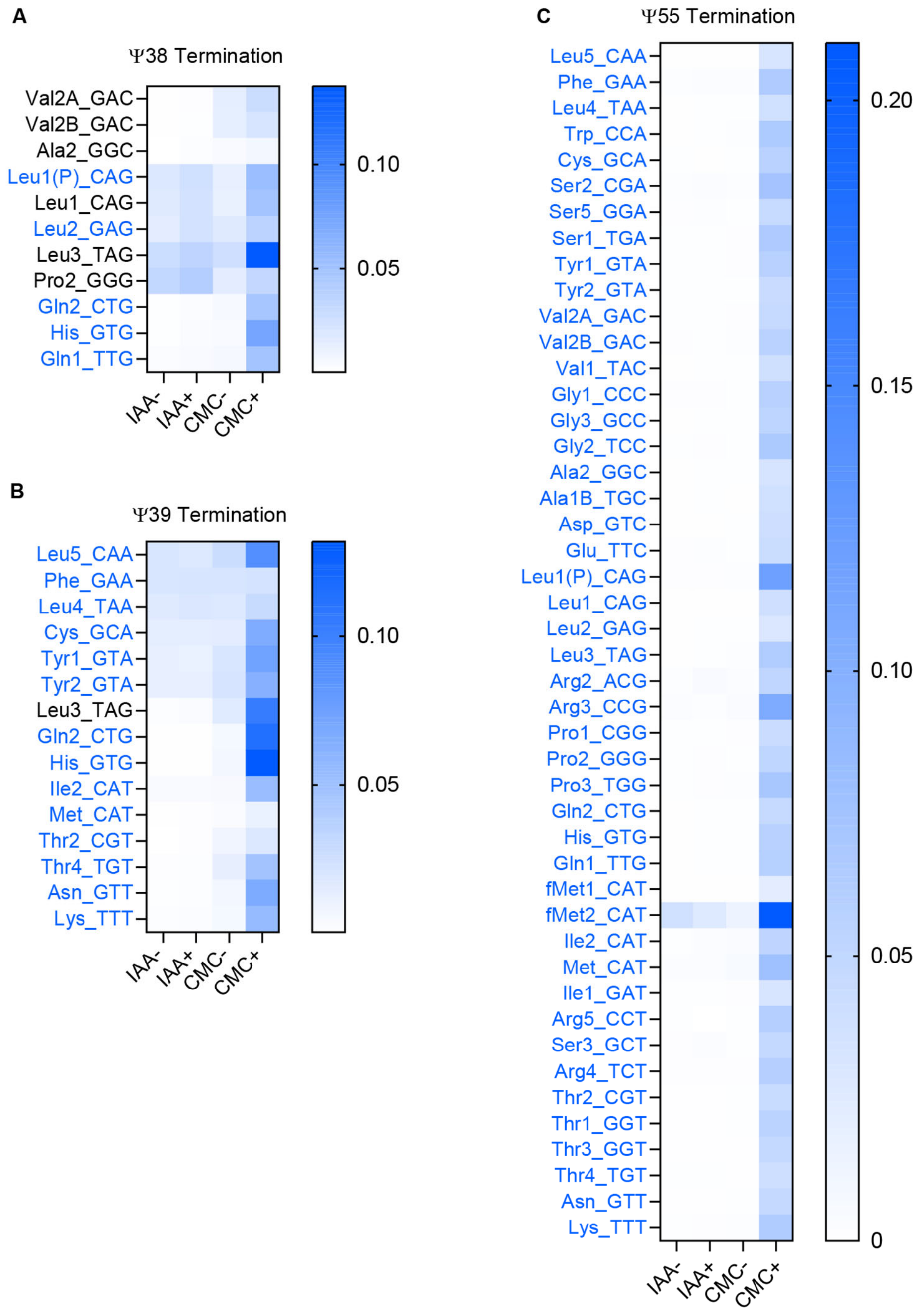


**Supplementary Fig. 4. CMC treatment facilitate detecting Ψ.**

Heatmaps of the termination frequencies at the uridine in *E. coli* tRNAs at position 40 (A), 41 (B), and 57 (Ψ55) (C). tRNA species where U at the indicated position is known to be Ψ are shown in blue. Chemical treatment is shown in bottom.

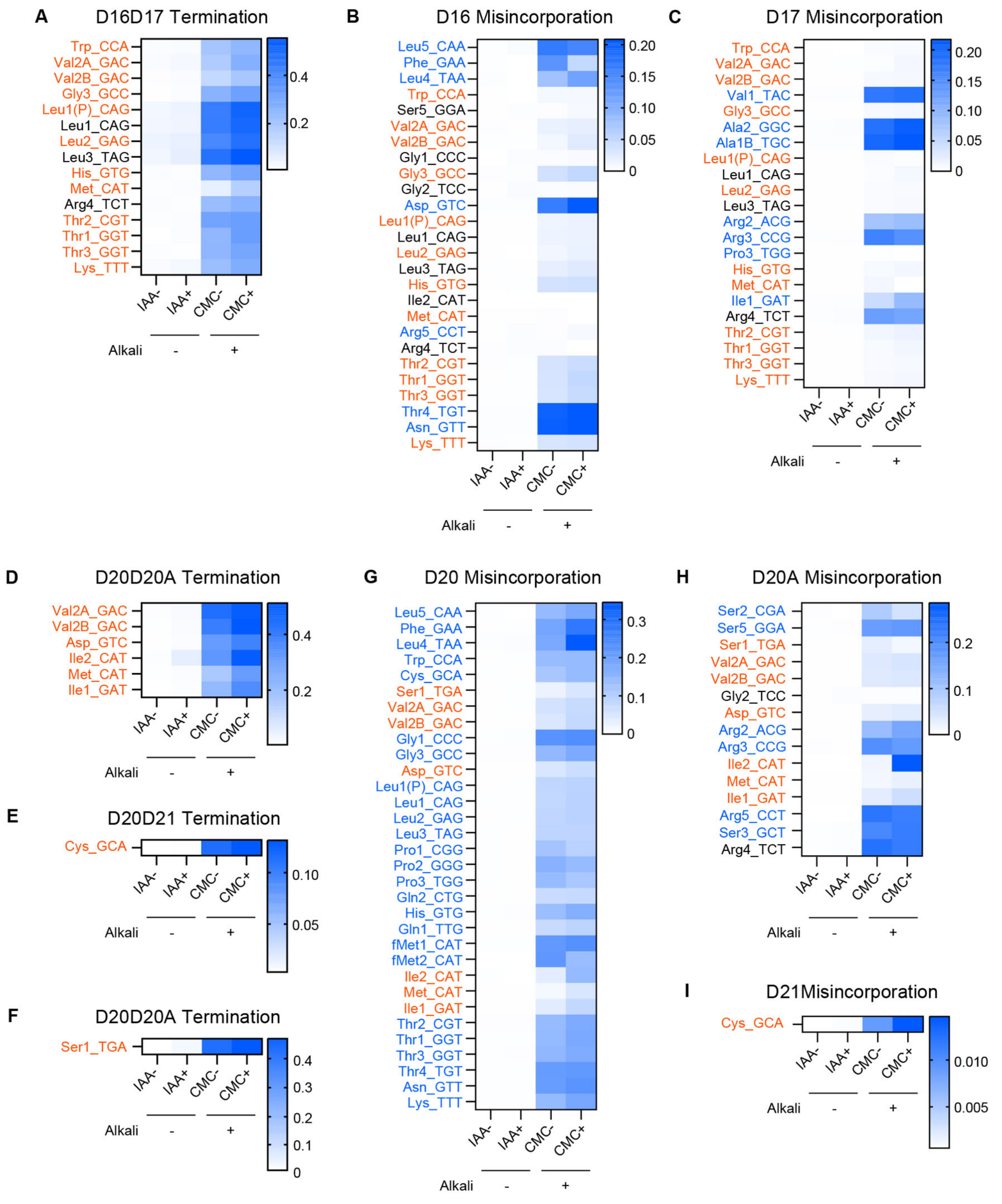


**Supplementary Fig 5. Misincorporation and termination signals derived from D in *E. coli* tRNAs.** Heatmaps showing termination frequencies at position 18 (A), 21 (D and E), and 22 (F) , and misincorporation frequencies at position 16 (B), 17 (C), 20 (G), 20A (H), and 21 (I) to predict the presence of D. tRNAs bearing U at the indicated positions are shown. tRNA species bearing two consecutive D at positions are shown in red, whereas tRNA species containing single D at positions are shown in blue. D at a single position appears to induce misincorporation, and consecutive D likely induces termination of reverse transcription at the following position. The RT-signatures are elevated in CMC- and CMC+ conditions, which include alkali treatment. Strain and chemical treatment are shown at the bottom.

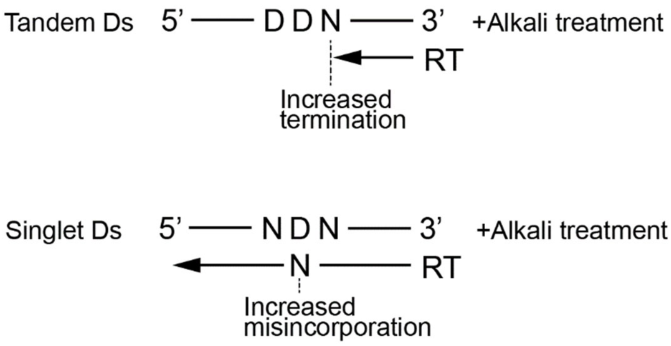


**Supplementary Fig 6. RT-derived signatures originated from tandem Ds or singlet Ds**

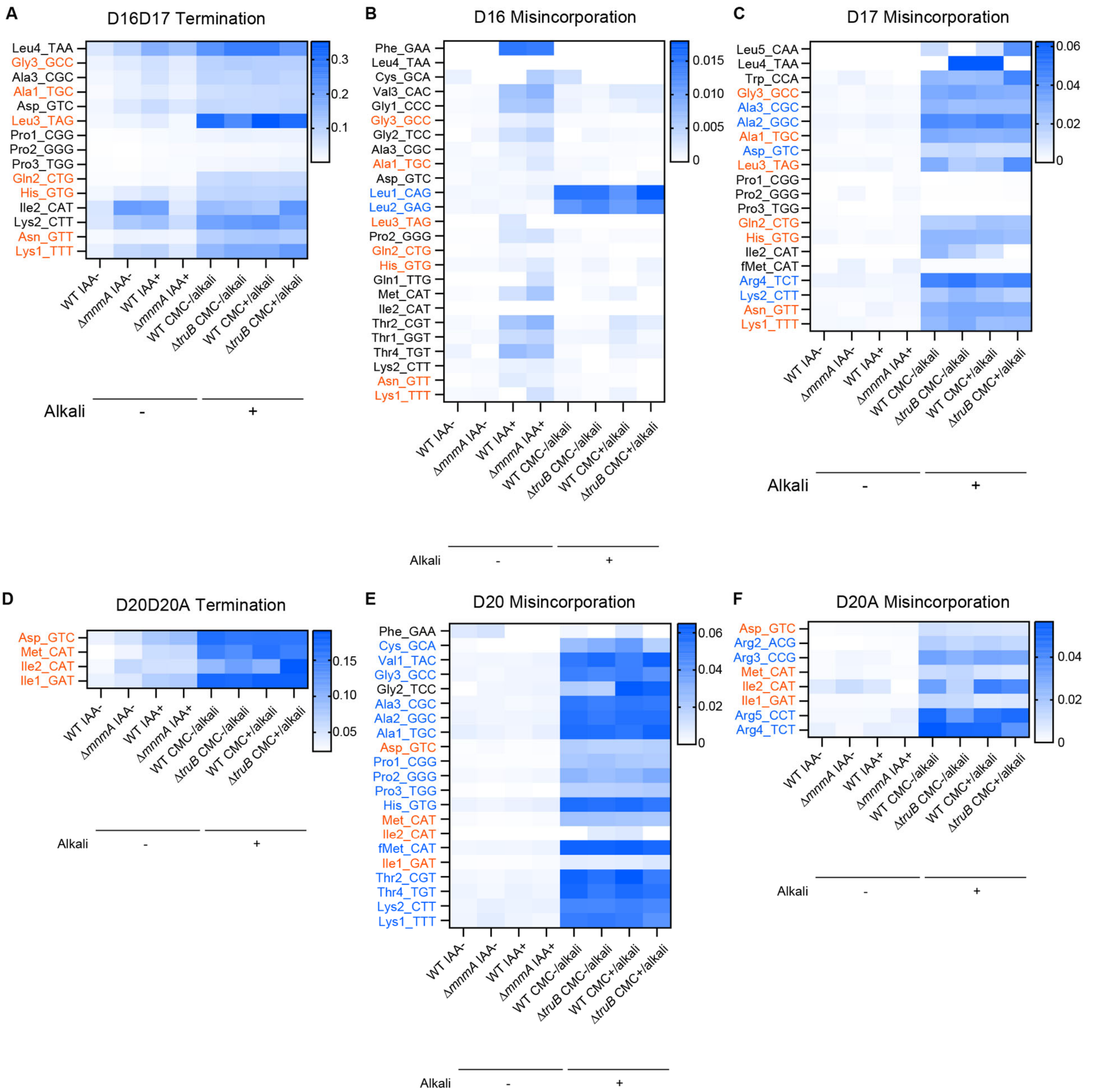


**Supplementary Fig. 7. Misincorporation and termination signals derived from D at positions 20, 20A and 21 in *Mtb* tRNAs.** Heatmaps showing termination frequencies at position 18 (A) and 21(D), and misincorporation frequencies at position 16 (B), 17 (C), 20 (E), and 20A (F) to predict the presence of D. tRNAs bearing U at the indicated positions are shown. tRNA species bearing two consecutive D are shown in red, whereas tRNA species containing single D are shown in blue. D at a single position appears to induce misincorporation, and consecutive D likely induces termination of reverse transcription at the following position. The RT-signatures are elevated in CMC- and CMC+ conditions, which include alkali treatment. Strain and chemical treatment are shown at the bottom.

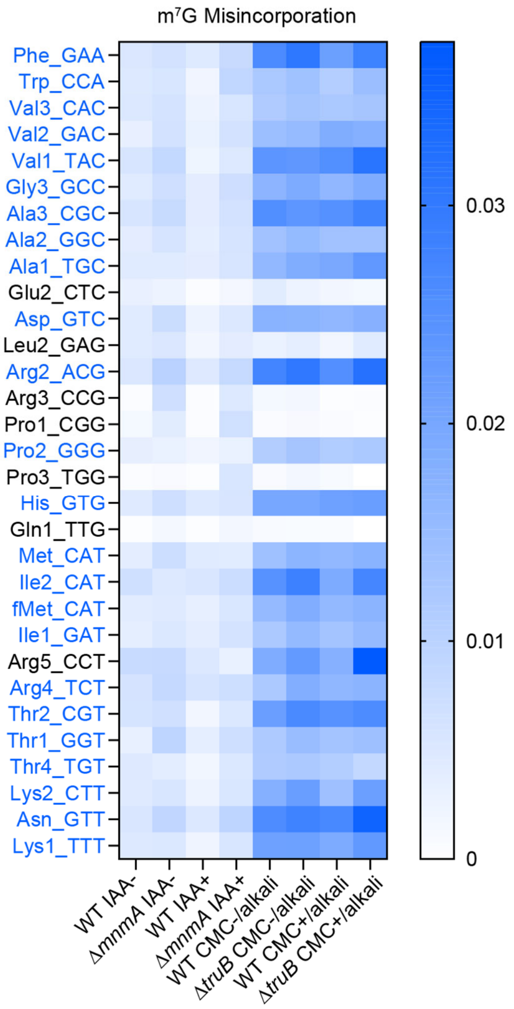


**Supplementary Fig. 8. Misincorporation signals derived from m$^7$G at position 46.** Heatmaps showing terhmianaion frequencies at position 46 to predict the presence of m$^7$G. tRNAs bearing G at position 46 are shown. tRNAs that have consistently high signals when tRNAs are treated alkali are shown in blue. Strain and chemical treatment are shown in bottom.

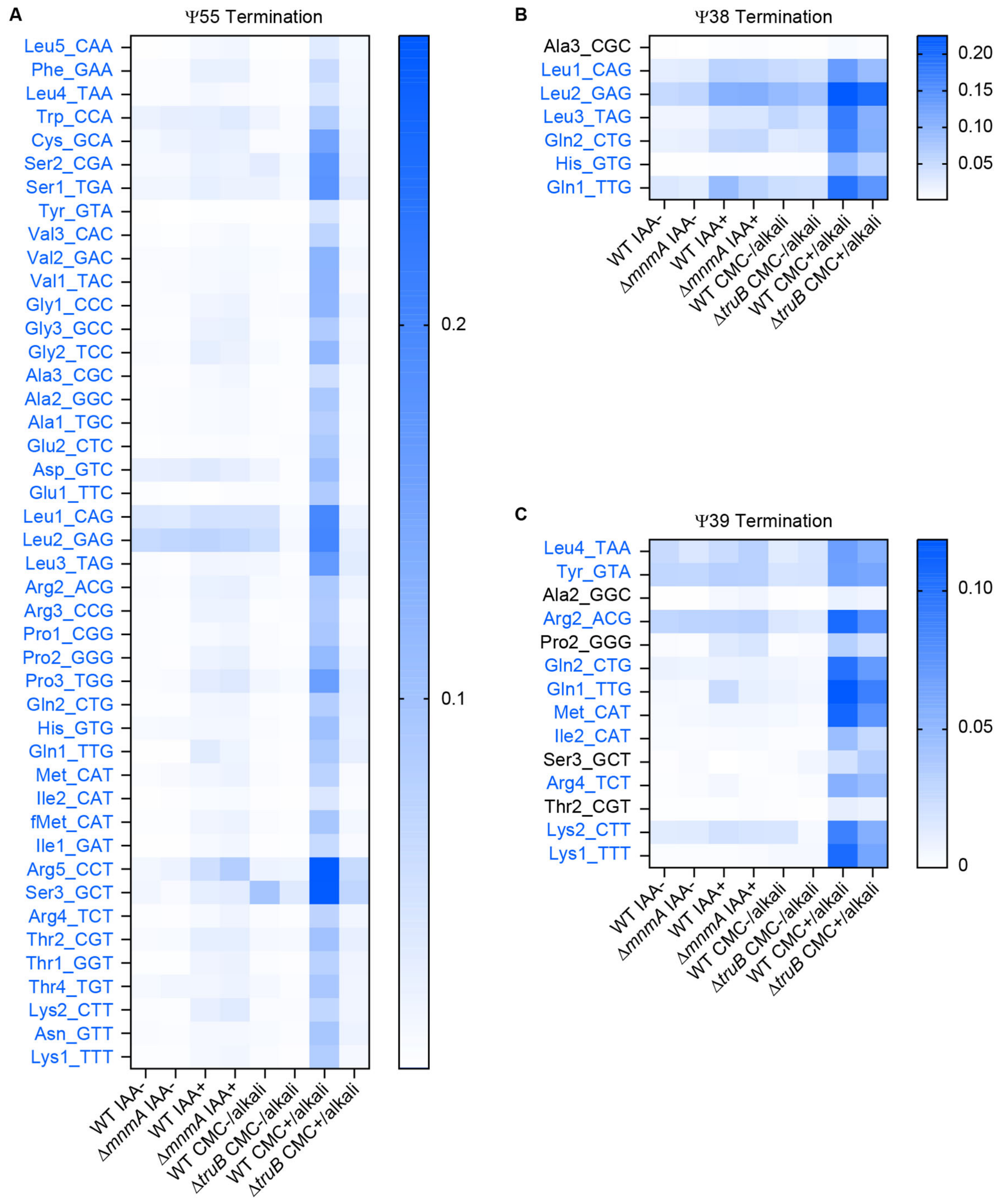


**Supplementary Fig. 9. Termination signals derived from pseudouridine at position 55, 38, and 39.** Heatmaps showing termination frequencies at position 57 (A), 40 (B), and 41(C) to assess the pseudouridylation states at position 55 (A), 38 (B), and 39 (C). tRNAs bearing U at the indicated positions are shown and tRNAs that have higher signals when CMC treated samples are shown in blue. Strain and chemical treatment are shown in bottom.
